# Domestic freezer storage: a solution for preserving stool microbiota integrity for at-home collection

**DOI:** 10.1101/2024.09.02.610874

**Authors:** Paula Momo Cabrera, Nicholas A. Bokulich, Petra Zimmermann

## Abstract

The gut microbiome is crucial for host health. Early childhood is a critical period for the development of a healthy gut microbiome, but it is particularly sensitive to external influences. Recent research has focused on using advanced techniques like shotgun metagenome sequencing to identify key microbial signatures and disruptions linked to disease. For accurate microbiome analysis, samples need to be collected and stored under specific conditions to preserve microbial integrity and composition, with -80°C storage considered the gold standard for stabilization.

This study investigates the effect of domestic freezer storage on the microbial composition of stool samples from 20 children under 4 years with the use of shotgun metagenomic sequencing. Fresh stool samples were aliquoted into sterile tubes, with one aliquot stored at 4°C and analyzed within 24 hours, while others were frozen in domestic freezers (below -18°C) and analyzed after 1 week, 2 months, and 6 months. Assessments of contig assembly quality, microbial diversity, and antimicrobial resistance genes revealed no significant degradation or variation in microbial composition.

**Importance:** Most previous studies on sample storage have used amplicon sequencing, which limits relevance to metagenome sequencing, in which contig quality and functional gene detection are additional concerns. Moreover, the effects of domestic freezer storage for at-home stool collection on microbiome profiles, contig quality, and antimicrobial resistance gene profiles have not been tested previously.

Our findings suggest that stool samples stored in domestic freezers for up to six months maintain the integrity of metagenomic data. These findings indicate that domestic freezer storage does not compromise the integrity or reproducibility of metagenomic data, offering a reliable and accessible alternative for temporary sample storage. This approach enhances the feasibility of large-scale at-home stool collection and citizen science projects, even those focused on the more easily perturbed early life microbiome. This advancement enables more inclusive research into the gut microbiome, enhancing our understanding of its role in human health.

## Introduction

The composition of the stool microbiome has been linked to many diseases, including allergic, inflammatory, rheumatological, metabolic, and psychiatric diseases(1–5). To accurately determine dysbiotic or suboptimal microbiome states and identify confounding factors, longitudinal sampling of large, representative population cohorts is essential. However, this can be logistically challenging. The current gold standard for preserving microbiome integrity is immediate DNA extraction or freezing of the stool sample at -80°C, as the addition of stabilisation buffers can affect DNA quantity and purity or lead to bacterial cell lysis(6, 7). While a number of studies have now tested refrigeration and room-temperature storage conditions(8–14), the impact of freezer temperature on microbiota composition during storage warrants further attention.

Immediate DNA extraction is not always feasible, particularly in studies with geographically dispersed participants. Some studies show that DNA integrity and microbial composition are maintained with storage of the stool at room temperature for up to 24 hours(8, 15), at 4°C for up to 24–72 hours(15–18), and at −20°C for up to 7 days(15, 18, 19). Conversely, other studies indicate that storage of the stool at room temperature over 12–24 hours(9, 16, 20) and at −20°C over 3–7 days(21) results in significant changes in bacterial composition. One study has even reported changes in the relative abundance of bacterial phyla after just 30 minutes of exposure to room temperature(21). Recent studies suggest that storage of stool sample at −20°C may suffice for short-term preservation(22), challenging the necessity of −80°C freezing. Thus, it remains uncertain whether the widely accepted gold standard of −80°C freezing is truly necessary, offering a possibility for investigating alternative cost-effective and accessible storage solutions.

A key concern with using domestic freezers is the occurrence of freeze-thaw cycles, particularly in frost-free freezers. These cycles cause periodic temperature fluctuations, which have been linked to changes in microbiome composition some studies(19), but not in others(18, 21). Most modern household refrigerator-freezers are equipped with automatic frost removal systems that operate in cycles lasting between 10 to 30 minutes(23, 24), typically occuring once or twice daily. During these defrost cycles, the air temperature in the freezer can rise to approximately -4°C(25), which could potentially impact the stability and preservation of the microbial community.

In this study, we investigated the stability of microbial composition in stool samples from young children stored in home freezers (−18°C) over six months, a condition that has not been previously investigated and could pose significant logistical advantages to current sample collection and storage methods.

Enabling participants to store samples in their home freezers can reduce the frequency of nurse visits for sample collection, thereby lowering personnel and travel costs, as well as the logistical burden on participants and study coordinators. This becomes particularly relevant as the field moves towards larger cohort sizes and higher temporal density in microbiome samples.

## Results

### Stool microbial diversity

Principal component analysis on Aitchison distances **(Figure 1A)** indicates that inter-individual variation in microbial profiles significantly exceeded any differences attributable to storage duration. As illustrated in **Figure 1B**, alpha diversity remained stable over time, with no significant differences observed between time points, not even in comparison to the unfrozen samples at 0W. Additionally, the number of observed species **(Figure 1C)** showed no significant differences across time points. The mean Aitchison distance, which represents compositional differences between samples at different time points, showed no significant variations **(Figure 1D)**. Bray-Curtis and Jaccard distance metrics (**Figures S1A** and **S1B** respectively), further showed that inter-individual variability surpasses temporal variability.

**Figure 1.**
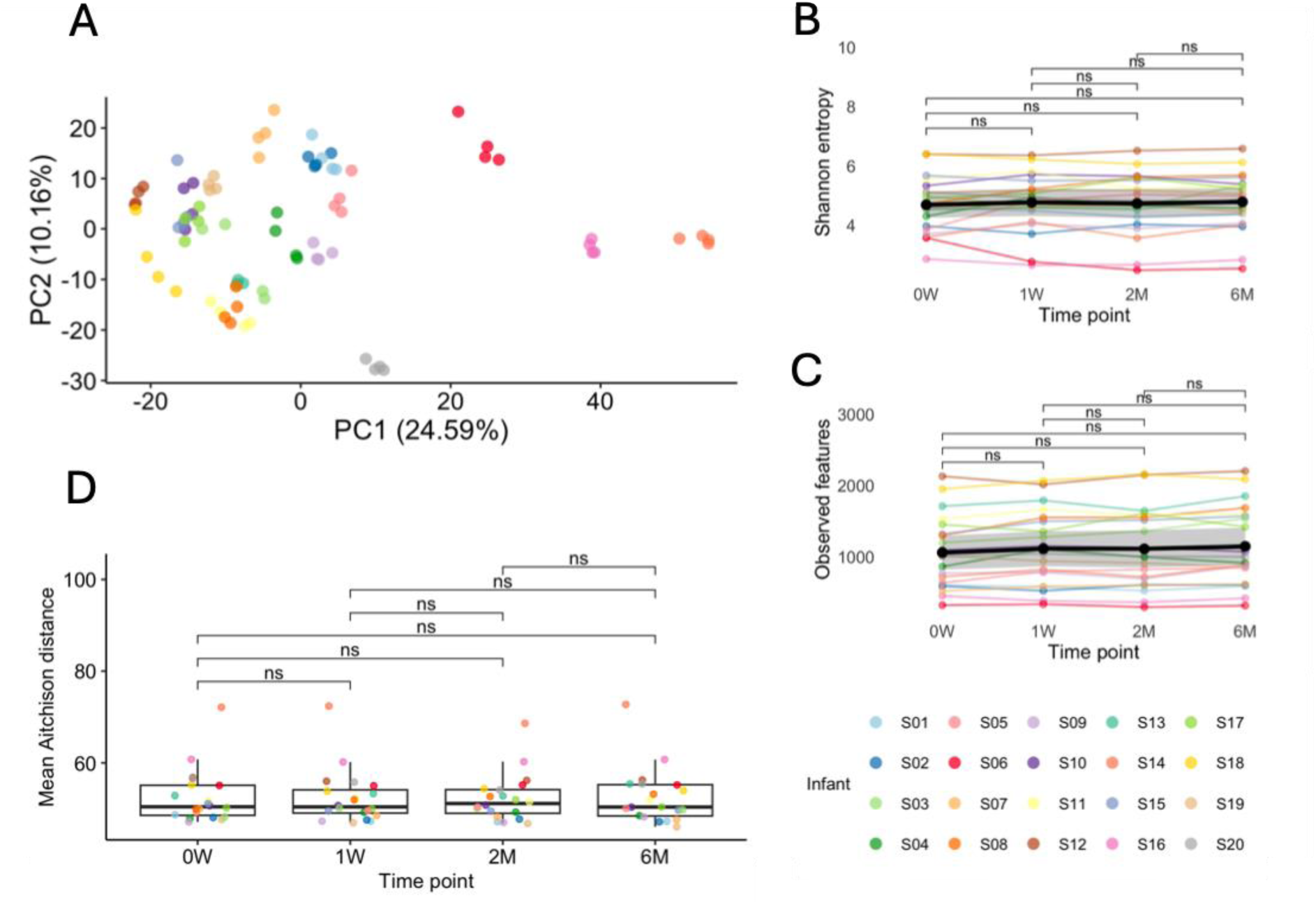
(A) PCoA based on the Aitchison distance matrix, illustrating the beta diversity of the microbial communities at different time points (0W = week 0, 1W = week 1, 2M = month 2, 6M = month 6). Each dot represents a sample, colored by child ID (S01-S20), indicating temporal shifts in microbial community composition. **(B)** Shannon diversity index over time, showing alpha diversity across different time points. **(C)** Number of observed features (species) over time, indicating the richness of the microbial communities. **(D)** Mean Aitchison distance between samples at different time points, representing compositional differences. The Wilcoxon rank-sum test with Bonferroni correction was used to assess differences, with no significant differences (ns) observed between the time points.

### Inter-individual differences have greater influence in stool microbial diversity than temporal effects

To better understand the factors influencing microbial community structure, we employed Linear Mixed Effects (LME) models **(Table S1)**. Our analysis revealed that storage time did not significantly affect microbial community composition when evaluated using Aitchison (p = 0.267, β = -0.03) and Jaccard (p = 0.836, β = 0.000) metrics, though a weak but significant effect was observed with Bray-Curtis (p = 0.007, β = -0.004).

In contrast, age emerged as a significant factor, consistently reflecting inter-individual variation in microbial communities. This was particularly evident in the Aitchison (p = 0.005, β = 3.062) and Jaccard (p = 0.004, β = 0.019) metrics, although the Bray-Curtis metric did not show a significant effect (p = 0.629, β = 0.006).

Further supporting the minimal impact of temporal changes, PERMANOVA results indicated no significant differences in beta diversity across different time points (p = 1 for Aitchison, p = 0.935 for Bray-Curtis, and p = 0.992 for Jaccard) **(Table S2)**.

To evaluate the relative contributions of temporal versus inter-individual variation in stool microbiome profiles, we implemented a Random Forest classifier to predict the sampling time points. The classifier’s performance was suboptimal, as evidenced by the confusion matrix **(Figure S2A)**, where no clear pattern of accurate classification emerged. The ROC curves **(Figure S2B)** further illustrate the classifier’s inefficacy, with area under the curve (AUC) values across different time points ranging from 0.10 to 0.24, only marginally above the expected performance of random guessing. Notably, the overall accuracy failed to exceed random chance, emphasizing the difficulty in distinguishing samples based solely on their microbiome profiles over time **(Figure S2C)**.

### Temporal dynamics of specific taxa

A total of 115 species were detected at or above 1% relative abundance; however, none showed significant deviations from the pre-freezing baseline (0W) following ANCOM-BC analysis. As can be osberved in **Figure 2**, subtle qualitative fluctuations were noted among subjects over time, these changes in the microbial community did not reach statistical significance neither at genus or species level, and are likely attributable to inherent sample heterogeneity rather than storage conditions.

**Figure 2.**
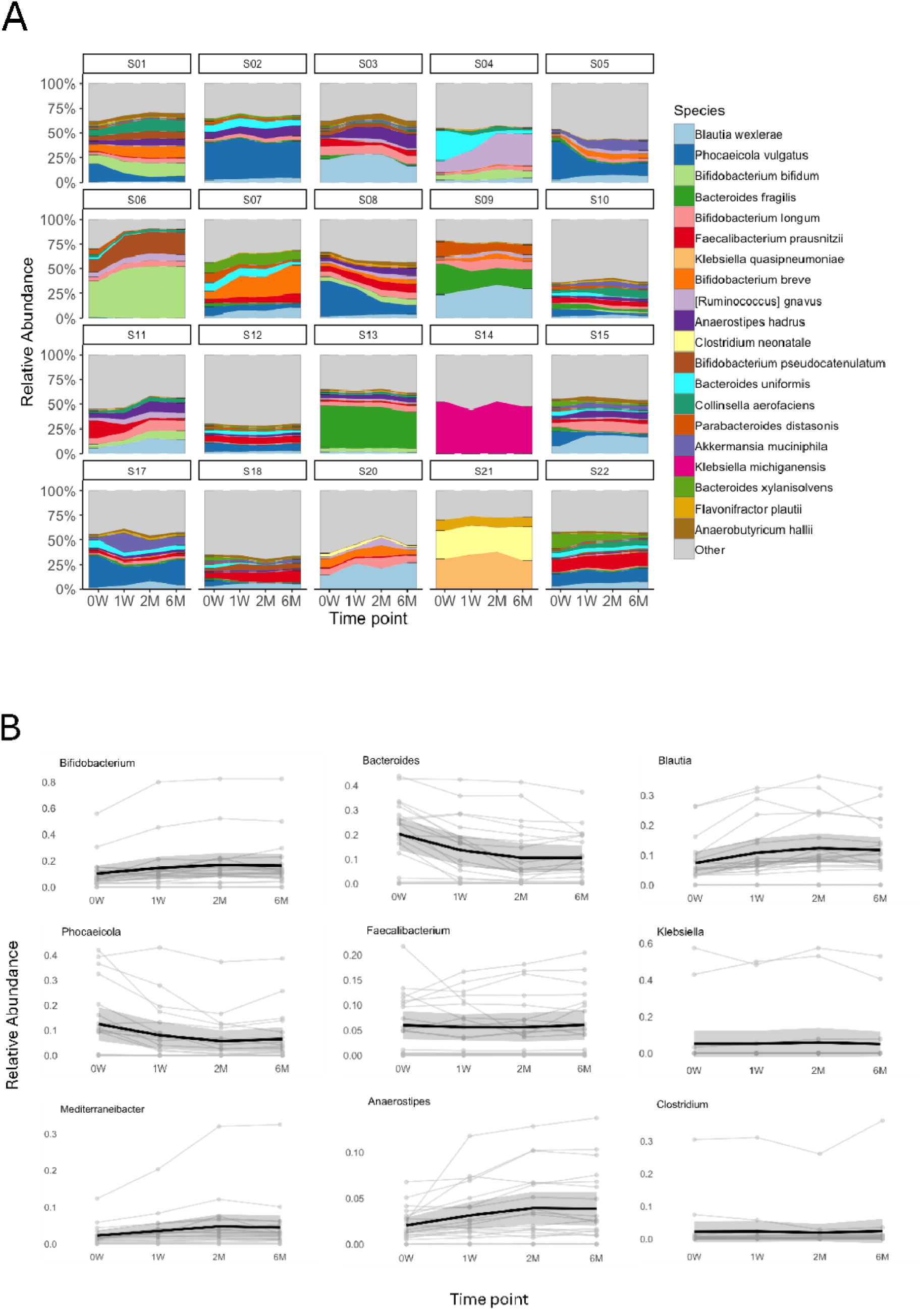
(A) Stacked area plots showing the relative abundance of the top 20 species across different time points (0W = week 0, 1W = week 1, 2M = month 2, 6M = month 6) for each child (S01–S20). Each colour represents a different species, illustrating the dynamic changes and stability in the microbial community composition over the first six months of life. **(B)** Line plots depicting the relative abundance trends of the most abundant genera. Each plot shows the relative abundance over time per individual child data (thin lines) and the overall mean trend (thick line). These plots highlight the temporal dynamics and individual variability within the microbial communities at the genus level.

### Temporal variability in antimicrobial resistance

In **Figure 3A**, the PCoA based on the Aitchison distance shows variability in AMR profiles across different subjects and time points. The first principal component (PC1) accounts for 25.82% of the variance, while the second principal component (PC2) explains 11.13%. To further understand the drivers of these observed changes, we analyzed the metadata variables contributing to the principal coordinates from the PCoA of Jaccard distance **(Figure S3)**. Specifically, time point (week) (R² = 0.04 for PC1) and “age” (R² = 0.013 for PC2) demonstrated to be poorly correlated, consistently revealing the lack of impact of sample collection time point in AMR profiles. Conversaly, child age (months) showed to be most substantial contributior to the observed variability (R² = 0.54 for PC1 and R² = 0.32).

**Figure 3.**
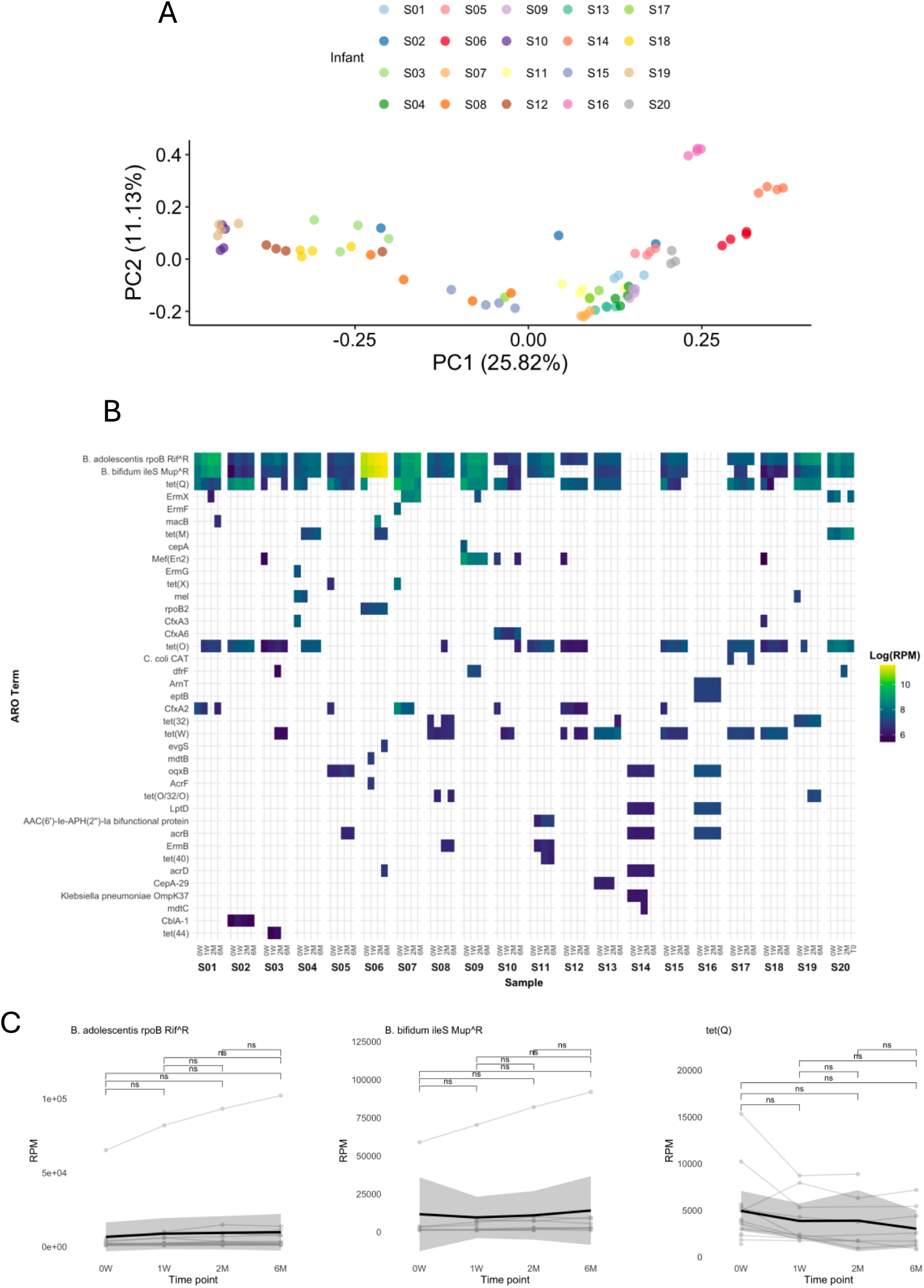
(A) Principal Coordinates Analysis (PCoA) plot depicting beta diversity based on the Aitchison distance matrix. Each point represents a sample, colored by child (S01-S20), as indicated in the legend, with the percentage of variance explained by PC1 and PC2. **(B)** Heatmap of the top 20 most abundant ARO terms, displaying log-transformed normalized Reads Per Million (RPM) values **(C)** Line plots depicting the relative abundance trends of the most abundant amr-confering genes. Each plot shows RPM over time per individual child data (thin lines) and the overall mean trend (thick line). The Wilcoxon rank-sum test with Bonferroni correction was used to assess differences, with no significant differences (ns) observed between the time points.

Furthermore, **Figure 3B** displays antimicrobial resistance profiles over time. The analysis highlighted that the most prevalent AMR-conferring genes detected, including “B. adolescentis rpoB Rif^R” (mutated RNA polymyerase β subunit (*rpoB*) conferring resistance to rifampicin), and “B. bifidum ileS Mup^R” (mutated isoleucyl-tRNA synthetase gene (*ileS*), conferring resistance to mupirocin) showed stable mean abundance overtime across all children. I contrast, “tet(Q)” (tetracycline-resistant ribosomal protection protein gene) showed a trend of decreasing abundance over time **(Figure 3C).** However, none of this fluctuations resulted to be statistically significant **(Figure 3C)**.

### Contig assembly quality over time

Next, we evaluated the stability of contig assembly quality in stool microbiome samples over time to assess whether domestic freezer storage could induce DNA damage or other changes impacting assembly quality. We assessed key metrics such as N50 and L50 across four different time points: week 0 (0W), week 1 (1W), month 2 (2M), and month 6 (6M). These metrics are critical indicators of the quality and completeness of genome assemblies obtained from metagenomic sequencing data, representing the length of the shortest contig at the 50% genome assembly threshold (N50) and the number of contigs whose lengths sum to 50% of the genome assembly (L50).

The median N50 values showes slight fluctuations across the different time points, with no significant differences observed (ns) between 0W, 1W, 2M, and 6M. **(Figure 4A)**. Similarly, the L50 values across the same time points also exhibited minimal variation, with no significant differences (ns) detected between the different storage durations. **(Figure 4B)**.

**Figure 4.**
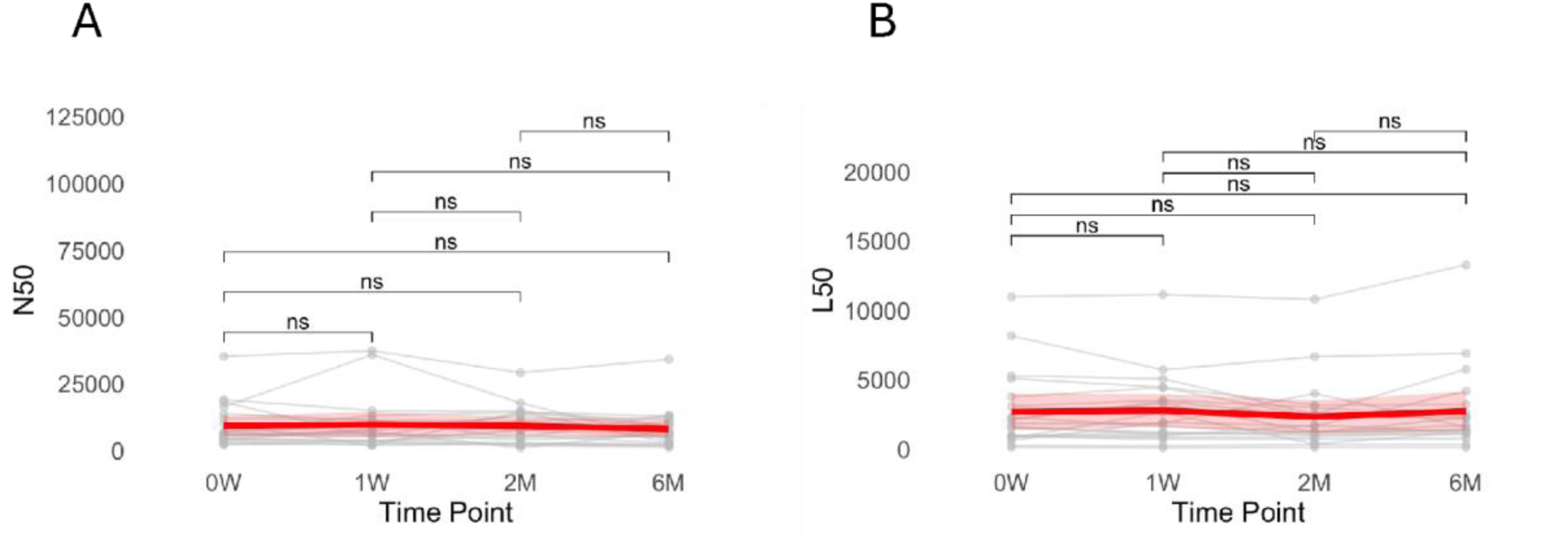
Longitudinal characterization of species-level relative abundance of 20 children’ stool microbiome samples of top 20 species over time. 0W= week 0; 1W= week 1; 2M= month 2; 6M= month 6. The Wilcoxon sum-rank test with Bonferroni correction was used to assess differences, with no significant differences (ns) observed between the time points.

## Discussion

In the rapidly evolving microbiome field, efforts have increasingly been directed towards larger cohort sizes and higher temporal density in sample collection. In this context, the need for reliable, cost-efficient, and low-burden sample collection and storage solutions have become paramount. Ensuring that storage conditions do not introduce significant biases or artifacts in metagenome sequence data is crucial for accurately tracking functional gene composition and microbial community dynamics over time.

Our findings demonstrate that DNA integrity, microbial and resistance-conferring gene diversity, as well as contig assembly quality, remain stable over a six-month storage period in home freezers. By employing gun metagenomic sequencing, we achieved species-level resolution in our analysis, allowing for a more comprehensive assessment of microbial diversity that extends beyond the conventional focus on taxonomic stability during storage. This provides a holistic view of stool DNA stability under storage in home freezers, offering valuable insights for optimizing long-term storage protocols. Additionally, the study’s design, which included multiple time points, adds robustness to our findings.

In this study, the core microbiome retains its richness and evenness over time, hence, irrespectively of the increasing number of freeze-thaw cycles endured during storage in home freezers. While contig assembly results were an essential quality check, our study used read-based analyses, such as microbial diversity and AMR profiling, which provides more detailed insights into the microbial community structure and function. While individual species’ and AMR relative abundances may fluctuate with storage in home freezers, the overall community structure remains largely stable. In fact, similarly to what was observed by Ilett et al.(26), our study found no significant differences between non-frozen 0W samples and the frozen 1W up to 6M samples. However, specific microbial taxa which were found to have subtle temporal fluctuations in abundance included *Bacteroides spp.* whose relative abundance decreased and *Bifidobacterium spp.* which increased over the six-month storage period. These taxa-dependent variations are consistent with previous studies and could be attributed to differences in bacterial cell wall structure, metabolic activity, and resilience to freezing and thawing processes(22, 27–29). These findings highlighted the intricate dynamics within microbial communities and the importance of considering taxa-specific responses when interpreting microbiome data. Our study identified both high stability in the most abundant and frequent resistance-conferring genes, and subtle shifts in those less abundant and highly infrequent. These shifts are likely more related to the heterogeneity of the samples than to storage conditions, emphasizing the complex and dynamic nature of microbial communities. The prevalence of highly infrequent and sparse AMR-confering genes suggest that while some resistance-confering genes maintain a consistent presence, others appear to be more sporadic and sparse. These findings aligned with previous reserach(27), which demonstrated that different storage methods could influence the detection and quantification of AMR-conferring genes, impacting the interpretation of resistance profiles in microbial communities and underscored the importance of longitudinal monitoring of AMR profiles to understand the evolution of resistance genes.

Our study addressed a significant gap in the literature by demonstrating the feasibility of using home freezer storage for preserving stool samples in home freezers. While different studies have investigated the temporal variability of a microbial community under different storage conditions, validating the stability of stool DNA stored in home freezers offers significant advantages for large-scale cohort studies, particularly those involving long-term follow-up and geographically dispersed participants. Notably, access to ultra-low temperature storage facilities (-80°C) is limited in many regions. By reducing the need for specialized storage facilities, researchers can optimize resource allocation and reduce logistical burdens on participants and study coordinators. This finding is particularly relevant for large-scale and citizen science projects, where cost-effective and accessible storage solutions are essential to promote inclusivity and broader participation in research(7).

Nevertheless, our study has several limitations. First, our study focused on a six-month storage period. Future research should explore longer storage durations to assess longer-term storage effects. Additionally, our study included a relatively small sample size of 20 childen, which may limit the generalizability of the results. A larger sample size would provide more robust data and allow for a more comprehensive analysis of variability across different individuals. Future research should also test the storage viability of a broader range of sample types to enable more extensive microbiome cohort studies and applications.

## Conclusion

The finding that home freezers can be used to effectively store stool samples for microbiome analysis significantly enhances the feasibility of long-term studies that involve at-home collection of stool samples. This approach promotes broader participation by allowing participants to conveniently store samples at home. Our results support the use of home freezer storage as a viable alternative to conventional methods, ensuring reliable and reproducible results. This advancement facilitates robust research into microbial dynamics and disease mechanisms. Future research should further explore the long-term stability of samples under various conditions to optimize preservation protocols and advance microbiome research.

## Materials and Methods

### Stool sample collection and storage in home freezers

Fresh stool samples from 20 healthy Swiss children (55% female) with a mean age of 22.4 (range 2 to 56) months, were aliquoted into 4 Fecon sterile tubes (Fecotainers, Medical Wire & Equipment Co. Ltd, United Kingdom). One aliquot was stored at 4°C and extracted and analyzed within 24 hours (0W), while the other three aliquots of the same sample were frozen in domestic freezers (below -18°C) and analyzed after 1 week (1W), 2 months (2M) and 6 months (6M). Samples were kept under the same conditions during transport to the Microbiota and Children Laboratory at the University of Fribourg, Switzerland.

### DNA extraction and metagenomic shot-gun sequencing

Aliquots of stool samples were thawed at room temperature, and 100 mg of each sample was used for DNA extraction using the FastDNA™ SPIN Kit for Soil (MP Biomedicals, Illkirch-Graffenstaden, France) according to the manufacturer’s instructions. DNA concentrations were measured using Qubit™ dsDNA High Sensitivity Assay kits (Life Technologies, California, United States). The DNA in the negative control was below the detection limit (<0.01 ng/µl) and was not sequenced. Sequencing libraries with an insert size of approximately 600 bp were prepared using Nextera DNA Flex library preparation kits (Illumina, San Francisco, United States), with the addition of Illumina PhiX DNA. Paired-end sequencing (2 × 149 bp) was performed on a NextSeq 550 system (Illumina) using high-output flow cells. Positive controls included bacterial and fungal mock communities (Gut Microbiome Whole Cell Mix MSA-2006™ and Mycobiome Whole Cell Mix, respectively, ATCC, Manassas, United States).

## Bioinformatic analyses

### Quality filtering and removal of human-mapping reads

FASTQC (v.0.11.9) was used to assess the quality of raw reads(30). Low-quality reads were filtered and trimmed using Trimmomatic v0.39(31) with the following settings: LEADING:3, TRAILING:3, SLIDINGWINDOW:4:20, MINLEN:36. Processed reads were then imported into QIIME 2 (Quantitative Insights Into Microbial Ecology 2)(32), where all further processing was performed using a QIIME 2 plugin for shotgun metagenome analysis (https://github.com/bokulich-lab/q2-moshpit). Bowtie2(33) and SAMtools(34) were used to align the reads against the human genome reference consortium (GRCh38)(35) to remove host sequences and retain non-host (unmapped) reads for subsequent analysis which resulted in a sum of 27,703,760 filtered reads across the dataset.

### Read-based microbiome taxonomy, diversity and antimicrobial resistance genes profiling

The high quality and filtered reads were taxonomically classified using Kraken2(36) using the Standard database. This was followed by abundance re-estimation using Bracken(37) to enhance taxonomic profiling accuracy. Rarefied reads subsampled to the minimum sample read depth, in this case, 346,297 taxonomically annotated sequences per sample (excluding the unclassified portion), were compared on species-level dissimilarity based on beta diversity metrics, including Aitchison, Bray- Curtis and Jaccard distances computed in QIIME 2. Additionally, alpha diversity metrics such as Shannon diversity and observed features (community richness) were also computed in QIIME 2.

To assess antimicrobial resistance (AMR) potential, reads were annotated using the Comprehensive Antibiotic Resistance Database (CARD)(38) in QIIME 2 using the q2- amr plugin (https://github.com/bokulich-lab/q2-amr). This involved using the Resistance Gene Identifier (RGI) application to predict antibiotic resistomes from protein or nucleotide data based on homology and single nucleotide polymorphsms (SNP) models.

### Metagenomic assembly and contig quality assessment

Trimmed and filtered reads from the individual samples were assembled into contigs using MEGAHIT(39) with default parameters using the QIIME 2 q2-assembly plugin (https://github.com/bokulich-lab/q2-assembly). Contig quality was assessed using metaQUAST(40) to calculate basic statistics such as assembly length, N50 values, and L50 metrics.

### Statistical analysis

Principal Coordinate Analysis (PCoA) was conducted using Aitchison, Bray-Curtis, and Jaccard distances in QIIME 2. To compare the mean Aitchison dissimilarity between different time points, the Wilcoxon rank-sum test was applied using the vegan package (v2.6-4) in RStudio (v2024.04.0+735), with p-values adjusted for multiple comparisons using the Bonferroni method. Additionally, PERMANOVA was performed with the vegan package (v2.6-4), utilizing the default number of permutations (n=999). Pearson correlations were employed to assess the relationship between numerical or transformed metadata variables and principal components (PC1 and PC2) derived from the distance matrices. The Wilcoxon rank-sum test with Bonferroni correction was also used to evaluate differences in alpha diversity metrics, specifically Shannon diversity and observed features, across different time points. Differentially abundant taxa, with a minimum relative abundance of 1% across all samples, were identified at both species and genus levels using ANCOM-BC(41).

To assess the influence of temporal versus inter-individual variation, we used the q2- sample-classifier plugin(42) (https://github.com/qiime2/q2-sample-classifier) in QIIME 2 to train a Random Forest classifier to predict sample time points based on stool microbiome profiles. The analysis employed standard parameters, including an 80/20 split for training and testing and 100 trees in the Random Forest model, with performance evaluated through stratified k-fold cross-validation to determine the model’s accuracy in predicting sample time points.

Linear mixed-effects (LME) models were employed to assess temporal trends in the microbial community composition and diversity while accounting for repeated measures within the same subjects over different time points, and were performed using the QIIME 2 q2-longitudinal plugin(43) (https://github.com/qiime2/q2-longitudinal).

For constructing the heatmap for Antimicrobial Resistance Ontology (ARO) terms, normalized reads representing the number of sequence reads that were fully mapped to the reference sequence without any gaps were used to ensure accurate ARO term assignment. Mapped reads per sample were normalized for sequencing depth differences and to reduce sequencing bias by dividing the completely mapped reads by each sample’s total read count and multiplying by 1,000,000 to obtain Reads Per Million (RPM). These RPM values were then log-transformed to account for the variation in read ranges. The top 20 most abundant ARO terms were plotted in a heatmap using ggplot2 (v3.5.1) in RStudio (v2024.04.0+735).

Statistical analysis based on the Wilcoxon sum-rank test with Bonferroni correction of the N50 and L50 metrics was conducted to assess the quality of the metagenomic assemblies. These metrics were calculated and compared across the different storage time points to evaluate any changes in assembly quality.

## Data availability

The sequencing raw data is deposited on the European Nucleotide Archive (ENA) with accession PRJEB79382. Data will be made publicly available after publication. A STORMS (Strengthening The Organizing and Reporting of Microbiome Studies) checklist(44) is available at DOI:<u>10.5281/zenodo.13460391</u>.

## Ethical compliance

This study has been approved by the Commission cantonale d’éthique de la recherche sur l’être humain (The Cantonal Commission for Ethics in Research on Human Beings (CER-VD)) of the canton of Vaud, Switzerland (#2019–01567).

## Acknowlegments

We would like to acknowledge Silvio Ghezzi for fecal sample collection and transport, and William Jakob for shotgun metagenomic sequencing data generation. We thank Dr. Michal Ziemski for his support on shotgun metagenomic sequencing analysis, and Dr. Anton Lavrienko for his valuable comments on the manuscript. This research was by two grants from the Swiss National Science Foundation (Grant 10000835 and PZPGP3_193140).

## Supplementary material DNA concentration

The mean DNA concentration per sample was 44.99 ng/µL (range 1.77 – 60 ng/µL). When analyzed on a per-child basis across all time points, some children, such as child S08, exhibited higher mean DNA concentrations with lower temporal variability, while others, like child S03, showed lower mean concentrations with higher temporal variability. Specifically, child S08 had a mean DNA concentration of 58.5 ng/µL (SD = 1.9 ng/µL), while child S03 displayed a mean concentration of 21.0 ng/µL (SD = 11.9 ng/µL), showing that individual children exhibited distinct patterns of DNA concentration measurements over time. However, Bonferroni-adjusted pairwise Wilcoxon tests among all children and time points revealed no significant differences in DNA concentrations.

**Figure S1.**
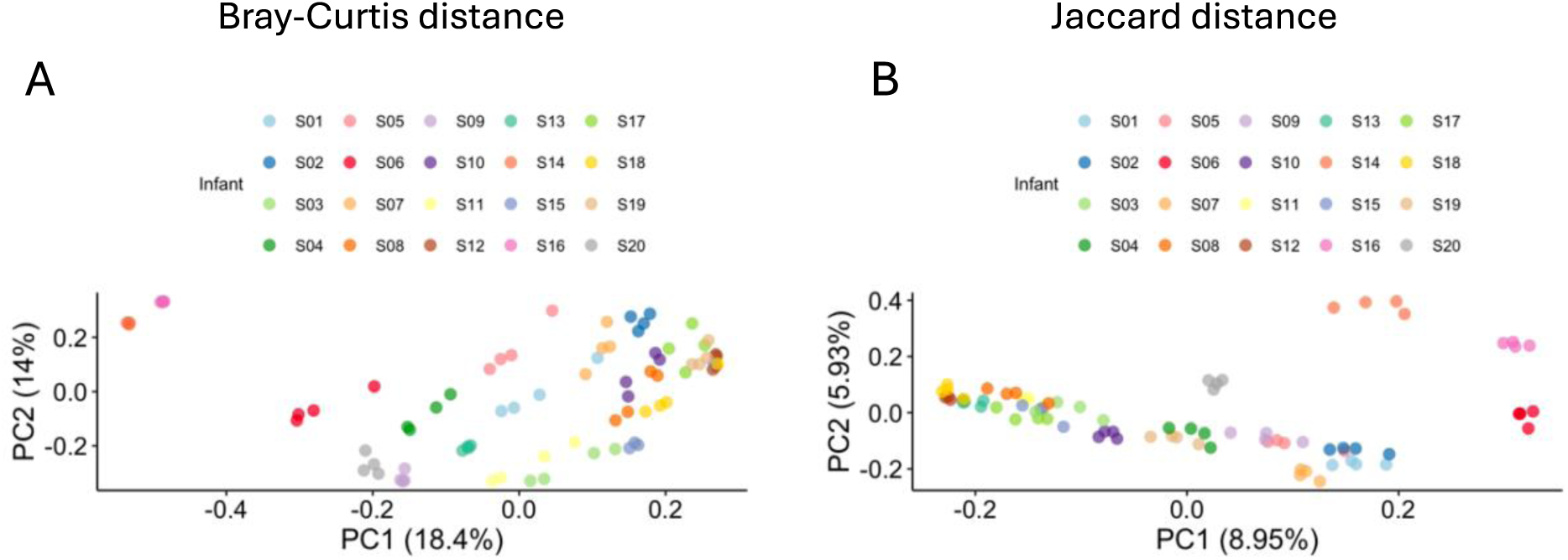
Principal Coordinates Analysis (PCoA) plot based on the **(A)** Bray-Curtis distance matrix and **(B)** Jaccard distance matrix illustrating the beta diversity of the microbial communities at different time points (0W = week 0, 1W = week 1, 2M = month 2, 6M = month 6).

**Figure S2.**
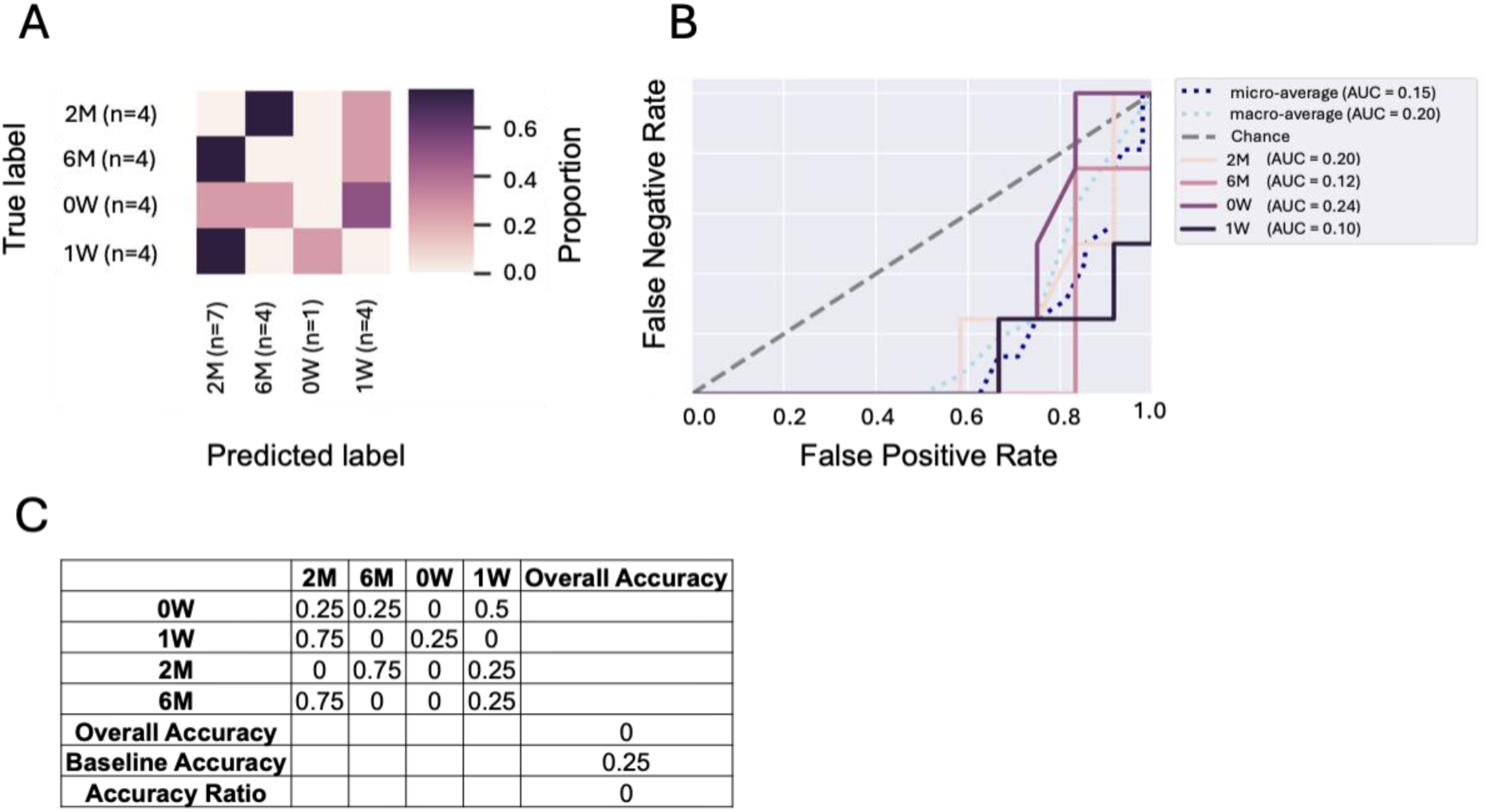
**(A)** Confusion matrix showing the Random Forest classifier’s performance in predicting time points (0W = week 0, 1W = week 1, 2M = month 2, 6M = month 6) based on microbiome data. Rows represent true time points, and columns represent predicted time points, with color intensity indicating the proportion of samples classified. **(B)** ROC curves for each time point with corresponding AUC values. The dashed line represents random chance, with low AUC values indicating poor differentiation between time points based on microbiome profiles. **(C)** Summary of the model accuracy results, including overall accuracy, baseline accuracy, and accuracy ratio, demonstrating the model’s performance across different time points.

**Figure S3.**
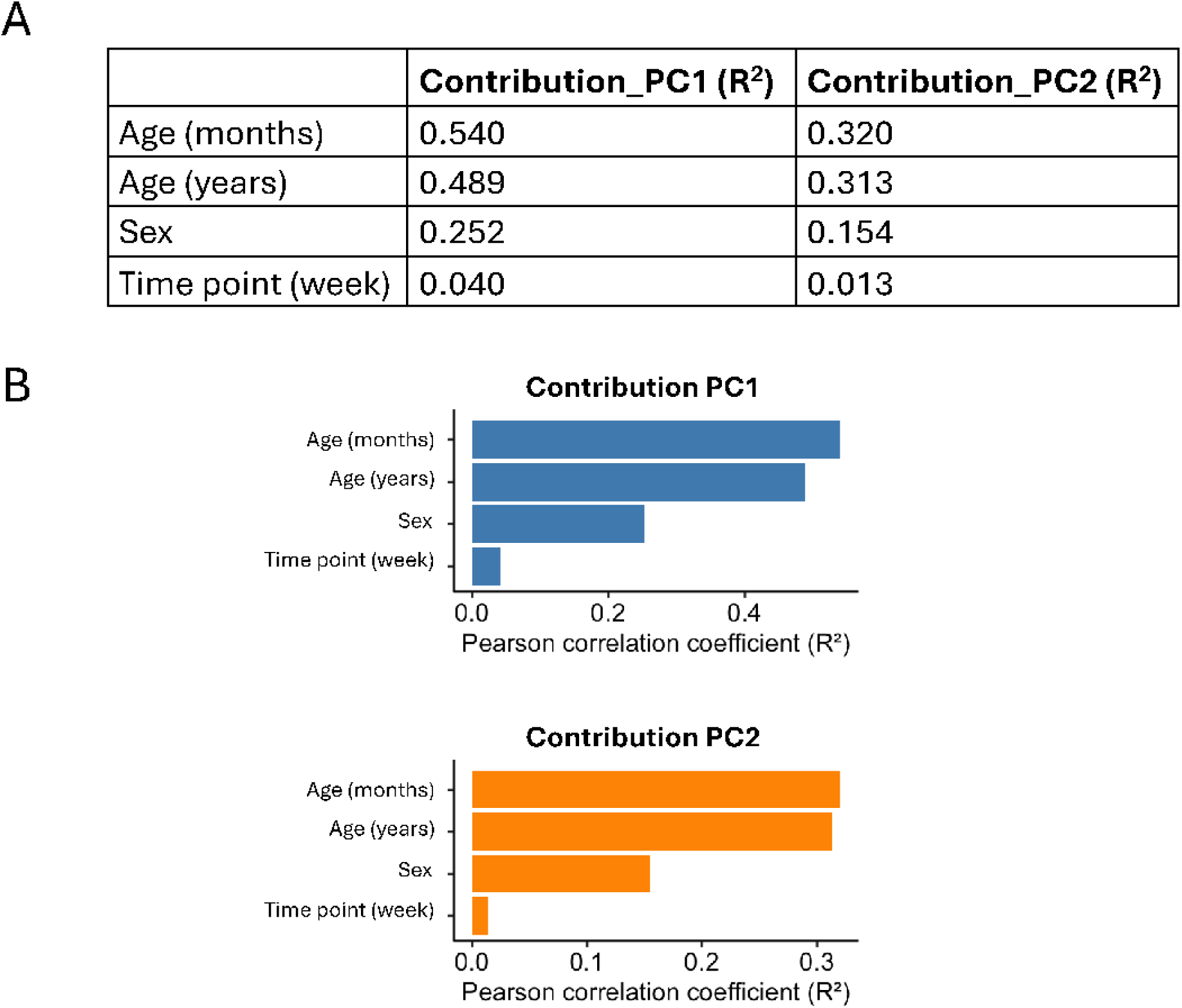
Metadata variables contributing to PC1 and PC2 from PCoA of Jaccard distance of the AMR profiles at all four time points. **(A)** Pearson correlation coefficients were calculated between numeric metadata variables and principal coordinates (PC1 and PC2) to assess their linear relationships. **(B)** The bar plots show the metadata variables with the highest absolute correlation to the first (PC1) and second (PC2) principal components. Contributions are calculated as the absolute values of the correlations between metadata variables and principal coordinates.

**Table S1.** Linear Mixed Effects (LME) Model results for beta diversity distance metrics: Aitchison Distance, Bray-Curtis Distance, and Jaccard Distance. Coef. (Coefficient (β)), Std. Err. (Standard Error), t-value (t-statistic value), P>|t| (p-value associated with the t-statistic), and [95% Conf. Interval] (95% Confidence Interval for the coefficient estimates).

Aitchison Distance
|  | Coef. | Std.Err. | z | P>\|z\| | [0.025 | 0.975] |
| --- | --- | --- | --- | --- | --- | --- |
| Intercept | 24.663 | 2.072 | 11.905 | 0 | 20.603 | 28.723 |
| Time point (week) | -0.03 | 0.027 | -1.111 | 0.267 | -0.083 | 0.023 |
| Age (year) | 3.062 | 1.092 | 2.804 | 0.005 | 0.922 | 5.202 |
| Time point:Age | 0.014 | 0.014 | 0.98 | 0.327 | -0.014 | 0.042 |

Bray-Curtis Distance
|  | Coef. | Std.Err. | z | P>\|z\| | [0.025 | 0.975] |
| --- | --- | --- | --- | --- | --- | --- |
| Intercept | 0.195 | 0.025 | 7.870 | 0.000 | 0.146 | 0.243 |
| Time point (week) | -0.004 | 0.002 | -2.677 | 0.007 | -0.008 | -0.001 |
| Age (year) | 0.006 | 0.013 | 0.483 | 0.629 | -0.019 | 0.032 |
| Time point:Age | 0.000 | 0.001 | 0.569 | 0.569 | -0.001 | 0.002 |

Jaccard Distance
|  | Coef. | Std.Err. | z | P>\|z\| | [0.025 | 0.975] |
| --- | --- | --- | --- | --- | --- | --- |
| Intercept | 0.644 | 0.012 | 51.900 | 0.000 | 0.620 | 0.668 |
| Time point (week) | 0.000 | 0.000 | 0.207 | 0.836 | -0.001 | 0.001 |
| Age (year) | 0.019 | 0.007 | 2.917 | 0.004 | 0.006 | 0.032 |
| Time point:Age | -0.000 | 0.000 | -1.410 | 0.159 | -0.001 | 0.000 |

**Table S2.** Permutational Multivariate Analysis of Variance (PERMANOVA) results for Aitchison, Bray-Curtis, and Jaccard distance matrices comparing the effects of week, subject, and age on microbial community composition. Df, Degrees of Freedom; SumOfSqs, Sum of Squares; R2, R-squared; F, F-statistic; Pr(>F), p-value; Perm, permutations.

| Distance | Variable | Df | SumOfSqs | R2 | F | Pr(>F) | Perm |
| --- | --- | --- | --- | --- | --- | --- | --- |
| Aitchison | Time point (week) | 1 | 1201.585 | 0.00656786 | 0.5156799 | 1 | 999 |
|  | Residual | 78 | 181747.721 | 0.99343214 | - | - | - |
|  | Total | 79 | 182949.306 | 1 | - | - | - |
| Aitchison | Infant | 19 | 124822.92 | 0.6822815 | 6.78139 | 0.001 | 999 |
|  | Residual | 60 | 58126.38 | 0.3177185 | - | - | - |
|  | Total | 79 | 182949.31 | 1 | - | - | - |
| Aitchison | Age (year) | 1 | 17349.92 | 0.0948346 | 8.172096 | 0.001 | 999 |
|  | Residual | 78 | 165599.38 | 0.9051654 | - | - | - |
|  | Total | 79 | 182949.31 | 1 | - | - | - |
| Bray-Curtis | Time point (week) | 1 | 0.1396905 | 0.00616375 | 0.4837546 | 0.935 | 999 |
|  | Residual | 78 | 22.5235331 | 0.99383625 | - | - | - |
|  | Total | 79 | 22.6632236 | 1 | - | - | - |
| Bray-Curtis | Infant | 19 | 21.022913 | 0.92762237 | 40.47291 | 0.001 | 999 |
|  | Residual | 60 | 1.640311 | 0.07237763 | - | - | - |
|  | Total | 79 | 22.663224 | 1 | - | - | - |
| Bray-Curtis | Age (year) | 1 | 1.73204 | 0.07642514 | 6.454442 | 0.001 | 999 |
|  | Residual | 78 | 20.93118 | 0.92357486 | - | - | - |
|  | Total | 79 | 22.66322 | 1 | - | - | - |
| Jaccard | Time point (week) | 1 | 0.1596754 | 0.00566108 | 0.4440781 | 0.992 | 999 |
|  | Residual | 78 | 28.046155 | 0.99433892 | - | - | - |
|  | Total | 79 | 28.2058304 | 1 | - | - | - |
| Jaccard | Infant | 19 | 24.401802 | 0.8651333 | 20.25703 | 0.001 | 999 |
|  | Residual | 60 | 3.804028 | 0.1348667 | NA | NA | - |
|  | Total | 79 | 28.20583 | 1 | NA | NA | - |
| Jaccard | Age (year) | 1 | 1.720459 | 0.06099656 | 5.066789 | 0.001 | 999 |
|  | Residual | 78 | 26.485372 | 0.93900344 | NA | NA | - |
|  | Total | 79 | 28.20583 | 1 | NA | NA | - |

**Table S3.** Permutational Multivariate Analysis of Variance (PERMANOVA) results for the Jaccard distance matrix comparing the effects of week, subject, and age on AMR profile. Df, Degrees of Freedom; SumOfSqs, Sum of Squares; R2, R-squared; F, F-statistic; Pr(>F), p-value; Perm, permutations.

| Distance | Variable | Df | SumOfSqs | R2 | F | Pr(>F) | Perm |
| --- | --- | --- | --- | --- | --- | --- | --- |
| Jaccard | Time point (week) | 1 | 0.1254 | 0.00718 | 0.5421 | 0.965 | 999 |
|  | Residual | 75 | 17.342 | 0.99282 | - | - | - |
|  | Total | 76 | 17.4674 | 1 | - | - | - |
| Jaccard | Infant | 19 | 9.4228 | 0.53945 | 3.5139 | 0.001 | 999 |
|  | Residual | 57 | 8.0446 | 0.46055 | - | - | - |
|  | Total | 76 | 17.4674 | 1 | - | - | - |
| Jaccard | Age (year) | 1 | 0.3239 | 0.01854 | 1.4169 | 0.116 | 999 |
|  | Residual | 75 | 17.1435 | 0.98146 | - | - | - |
|  | Total | 76 | 17.4674 | 1 | - | - | - |

